# Calling song maturity in two-spotted cricket, *Gryllus bimaculatus*: its patterns and a possible physical explanation

**DOI:** 10.1101/207712

**Authors:** Atsushi Miyashita

**Affiliations:** Department of Psychology and Neuroscience, Dalhousie University, 1355 Oxford Street, Halifax, Nova Scotia, Canada

**Keywords:** Cricket, Gryllus, Song, Calling, Bioacoustics, Frequency, Maturation

## Abstract

Males of *Gryllus bimaculatus* (two-spotted cricket) emit acoustic signals by stridulating two forewings. One of their songs, calling song, plays a role in attracting females to mate, yet the significance of each song trait in attracting conspecific females remains unclear. Among such traits, the relevance of frequency component (i.e. song pitch) has been underestimated, as orthopterans had long been believed incapable of song pitch recognition. However, recent literatures suggested that ears of orthopteran species are capable of frequency recognition as mammalians do. My previous report demonstrated that their calling song recorded from mature adults has constant and pure peak frequency component around 5.7 kHz, further supporting a biological significance of the frequency component. In this study, I tracked its change over time in sexual maturity phase (i.e. from early adult phase). 300 calling songs were recorded over time from a pool of 122 adult crickets, as it required large number of animals because the crickets rarely sing at very early adult stage. A maturation process of calling song was observed, where the peak frequency distributed lower and more variable frequency in early adult phase (e.g. mean peak value was 4.9 kHz on day 3), then it gradually increased and converged to 5.8 kHz with two weeks. The coefficient of variance also decreased over the process, showing minimum around day 20. Also, I found that the young crickets (supposed to emit immature song), emit perfectly tuned calling song with 5.8 kHz peak in helium-substituted (80% Helium and 20% Oxygen) environment. These findings suggest that the robustly regulated frequency of the calling song is acquired during the early-to-mid adult stage, and it may be associated with sexual maturity of males. Also, the helium-substitution experiment suggests that physical resistance from surrounding gas molecules negatively impacts stability of calling songs of young males, implying that a muscle development and/or forewing hardening may help song maturation. This study highlights a biological significance of the frequency component, such that females may adaptively select sexually mature males based on the song trait.

## Introduction

Acoustic signals are widely used for communication among members of the animal kingdom, including primates (Pfefferle and Fischer, 2006; Riede and Fitch, 1999), birds (Cardoso et al., 2008; Hall et al., 2013), frogs (Wilczynski et al., 1993), and arthropods Elliott and Koch, 1985; Katsuki and Suga, 1960; Miyashita et al., 2016a). Physical parameters that characterize acoustic signals are frequency (corresponds to pitch), amplitude (corresponds to volume), or more complex syllable patterns (Gray, 1997; Simmons and Ritchie, 1996; Zollinger et al., 2012). The frequency spectrum is highly unique to each species (Bennet-Clark, 1998; Rodríguez et al., 2015) even close species emit songs with comparatively different frequencies (Libersat et al., 1994). Frequency component distributes continuously to form a frequency spectrum. The frequency spectrum is associated with premating isolation (Balakrishnan and Sorenson, 2006; Honda-Sumi, 2005) in which females are only attracted by mating songs with a proper frequency and other songs are likely to be ignored.

Another example of biological significance of the frequency components is directional hearing. In crickets, a suite of mechanistic studies demonstrated that their auditory systems are capable of directional hearing, allowing them to locate the direction of a sound source (Boyd and Lewis, 1983; Lakes-Harlan and Scherberich, 2015; Michelsen et al., 1994). Several physical models are proposed for directional hearing, including amplitude, time, and phase differences between the left and right ears. Studies indicated a directionality in the response of the cricket auditory system between the left and right receptors, and that such directionality can be observed selectively around the dominant (major peak) frequency of cricket calling songs (Boyd and Lewis, 1983; Michelsen et al., 1994). Such a left/right imbalance of the auditory response is assumed to contribute to the directional hearing of cricket species.

In the two-spotted cricket *G. bimaculatus*, I previously reported that the dominant frequency of calling songs is strictly regulated, and is unaffected by male body size or resonator (mirror and harp regions on forewing) size (Miyashita et al., 2016a). Based on the idea of directional hearing, this trait would be adaptive for males because songs with inappropriate frequency may not transfer the information of the song emitter’s (i.e. males) position to conspecific females. This function is especially critical in mating contexts, where females being attracted by male calling song seek conspecific males based on the calling songs as a navigator.

Theoretically, the harmonic vibratory frequency of a membrane instrument (e.g. tympani) is affected by density, tension, and diameter (size) of the membrane (Fletcher, 1992). In a previous study, I revealed that the size (membrane area) of the two major sound sources of *G. bimaculatus* (harp and mirror region of the forewings) varies depending on whole body mass (Miyashita et al., 2016a), suggesting that the other factors (e.g. membrane tension, density) determine the constant frequency of the cricket songs. The trait also implied the biological significance of the frequency component of *G. bimaculatus* calling song.

In this study, I examined whether such unique and consistent frequency value of mature *G. bimaculatus* calling songs are age-dependent. The age-dependency of any song traits may be associated with maturation, learning, senescence, etc., and clarifying these associations would provide meaningful insights and new hypotheses to cricket ethology. In *G. bimaculatus*, adult male crickets begin to emit mating songs gradually from post-adult day 3, but it takes two weeks to reach the maximum activity (author’s personal observation). Therefore, it is necessary to set up large number of animals to successfully record the mating songs and acquire a robust conclusion especially for songs in early adult phase.

## Materials and Methods

### Crickets

Crickets were purchased from Tsukiyono-farm (Gunma, Japan) and reared at 28°C on a 12-h light and 12-h dark cycle as previously reported (Kochi et al., 2016; Miyashita et al., 2016b). Food and water were provided *ad libitum*. Crickets were isolated in plastic cups (1 cricket/cup) at final nymph instar, and the day of final molt (i.e. day 1) to adult stage was recorded. All the crickets were kept at 28°C. For this study, I purchased three batches from the company. For a quality control purpose, I checked survival rates of the three batches, and there was no difference in survival among these batches (Supplementary Fig. S1).

### Cricket song recordings

Cricket songs were recorded using a microphone (F-112, SONY, Tokyo, Japan) connected to a linear PCM recorder (PCM-D100, SONY, Tokyo, Japan). The sampling rate was 48,000 Hz, and the data were stored as uncompressed 16-bit wave files. Recordings were performed in a quiet room under white fluorescent light between January 23^rd^ and March 12^th^ in 2017, using crickets that emerged as adult on different dates. To record their songs, a sound recorder (turned on) was placed in front of multiple cricket-containing cups. The experimenter sat in the room and waited for crickets to sing. When a cricket sang, the cricket ID#, recording date, was recorded. In case two or more crickets sang, I waited for them to sing independently, after rattling their containers to make them silent. The cricket that has already been recorded its song on a day was then transferred to an incubator (so that the individual was no longer recorded on the day). Recording was performed daily (3-6 hours a day), and I successfully recorded calling songs from 63 out of 122 adult males (52%). The wave files were then processed to chunks of calling songs (typically 5 - 10 seconds) for later analyses. The calling songs from males no older than day 30 (post-adult) were used in the study, and the total number of recorded calling songs used in the analysis was 300. The number of recordings on each age (including those from males older than 30) is demonstrated in Supplementary Figure S2.

### Computing environment

Songs were analyzed on Macbook Pro (Retina, 15-inch, Mid 2015, OS X ver. 10.12.6, Apple Inc.) running R ver. 3.3.2, using the packages ‘seewave’ (ver. 2.0.5), ‘tuneR’ (ver. 1.3.2) for R. I also used Audacity version 2.1.1 (a sound analysis software on Mac OS X; http://audacityteam.org/) in this study.

### Calculation of peak frequencies of calling songs

Recorded wave files were loaded in R using the ‘seewave’ package, and a band-pass frequency filter (low-frequency cut-off, 0.5 kHz; high-frequency cut-off, 20 kHz) was applied. To obtain the peak frequency of the calling, I analyzed the spectrum of each calling song using the ‘spec’ function of the package, which returned frequency values and corresponding amplitude values. The Nadaraya–Watson kernel regression estimate was used to smooth the obtained data, and the peak frequency was determined as the frequency with the highest amplitude. This method returned peak values comparable with values obtained using Audacity (using ‘Plot Spectrum’ feature).

### Drawing figures

I used R ver. 3.3.2 to draw all figures shown in this paper. Although detailed information is provided in figure captions, it is important to note here that some longitudinal data are smoothened by applying moving median calculation to reduce fluctuating noises. Although moving median (or moving average) is commonly used for fluctuating time-series data to visualize general trends, I also included unsmoothed data in those figures to preserve data transparency. Also, a raw scatterplot of peak frequencies from 300 recordings was provided as a supplementary material (see Supplementary Fig. S3).

### Helium-substitution experiment

A mixture of 80% Helium and 20% Oxygen (Manyusha, Tokyo, Japan) was used in this study. Crickets were first placed in a plastic bag containing air (1 cricket/bag), and calling songs were recorded. The crickets were then moved to a plastic bag containing the Helium-Oxygen mixture (1 cricket/bag), and calling songs were recorded. In this study, crickets on day 5 (post-adult age) were used.

## Results

### Post-adult life history of *G. bimaculatus* used in this study

As shown in Figure 1A, the maximum lifespan (days were counted from the day of final molt) of *G. bimaculatus* used in this study was 68 days, while more than 60% of the crickets died within 30 days (Fig. 1A). Also, the number of recordings (which reflects both sexual activity of male crickets and their survival rate) peaked at day 4 (Fig. S2A), while it peaked around between day 10 and day 20 when normalized by survival rate (Fig. S2B). After the peak, there was an overall decreasing trend in number of recordings (both absolute numbers and normalized values) towards late adult phase (Fig. S2). This observation indicates that male sexual activity peaks around week 2 and 3 post-adult. Considering these results, I decided to track changes in calling songs within 30 days after final molt. In the following analysis, I included 300 calling songs recorded from 63 adult males. The hardening of forewing occurs at the very beginning after final molt, such that the forewings first appear (being moist and soft) on the day of final molt, but gradually dry up in 3 days (Fig. 1B). There was no apparent change after the hardening period (i.e. after day 3) throughout the experimental window (Fig. 1B), suggesting that any change in calling song patterns after day 3 may not be associated with general hardening of the forewings.

**Figure 1.**
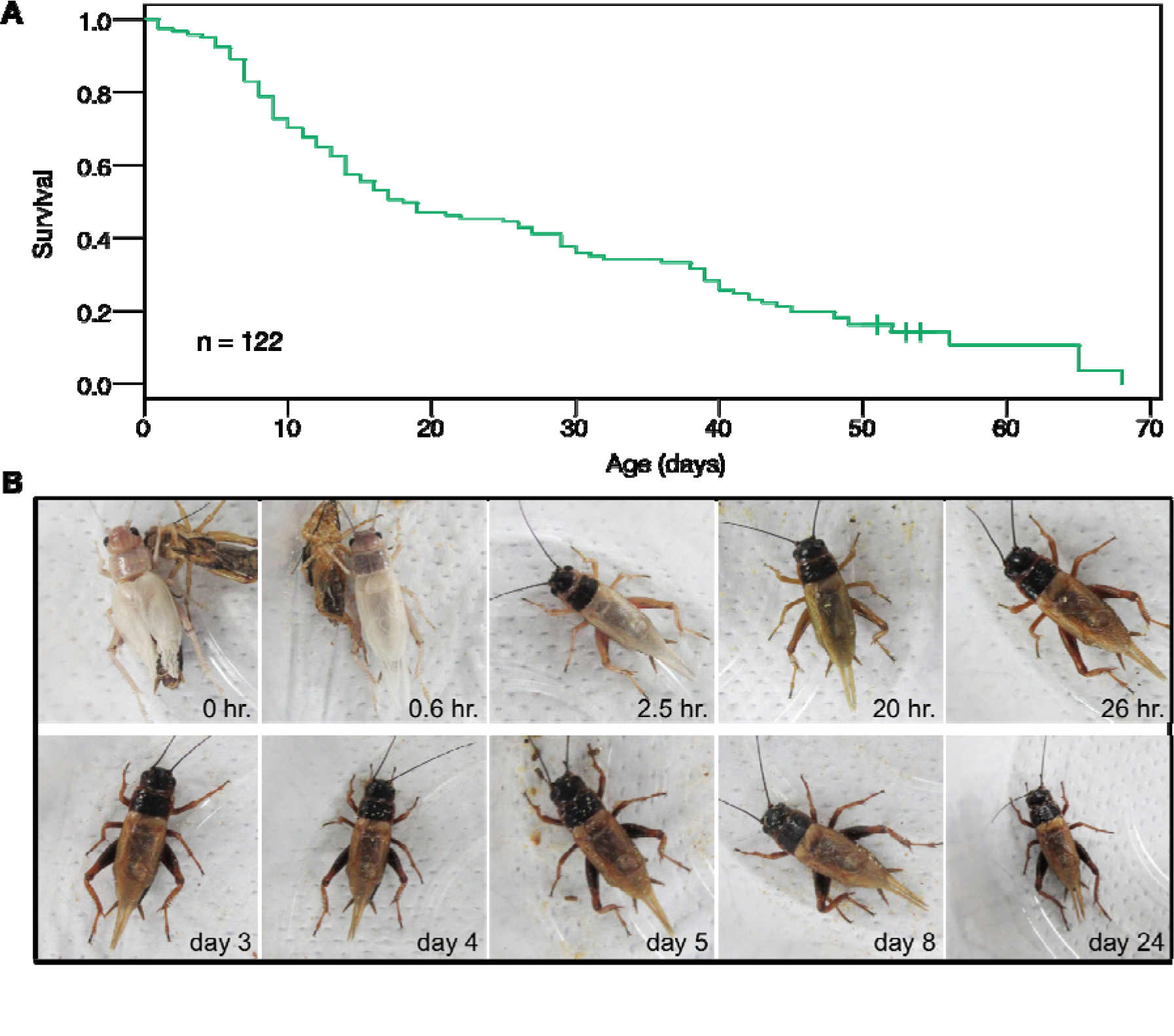
Survival and wing development of the crickets used in this study. A. Survival curve of the crickets (*G. bimaculatus*) used in this study. The horizontal axis shows male post-sdult age (counted from day of final molt), and the vertical axis shows survival. The green solid line shows Kaplan-Meier curve of the crickets. Ticks indicate that one or more crickets were censored at the time points (due to termination of experiment, or escaping). Green hatched area indicates 95% confidence intervals of the survival curve. This figure represents a pooled survival data of three independent batches of cricket (n=122). Survival curves separately drawn for each batch are shown in Supplementary Figure S1. B. Wing hardening of a male cricket after its final molt. Photos of a representative male *G. bimaculatus* were taken over time. The time at the lower right corner of each photo indicates the interval between final molt and when the photo was taken. The peak frequency of the calling song of this male was 4.9 kHz on day 4, and was 5.3 kHz on day 24.

### Age-dependent maturation of calling song

In this study, the crickets started emitting calling songs on day 2 at earliest (I obtained 1 recording on day 2, 8 recordings on day 3, 22 recordings on day 4, and 17 recordings on day 5 (Fig. S2A)). Calling songs recorded in the early period contained lower frequency peaks than those recorded in later adult phase (Fig. 2). The calling songs recorded on different ages from a representative male showed frequency peaks below 5.0kHz on day 4 and day 5, while it reached 5.8 kHz after day 8 and it remained (Fig. 2).

**Fig 2.**
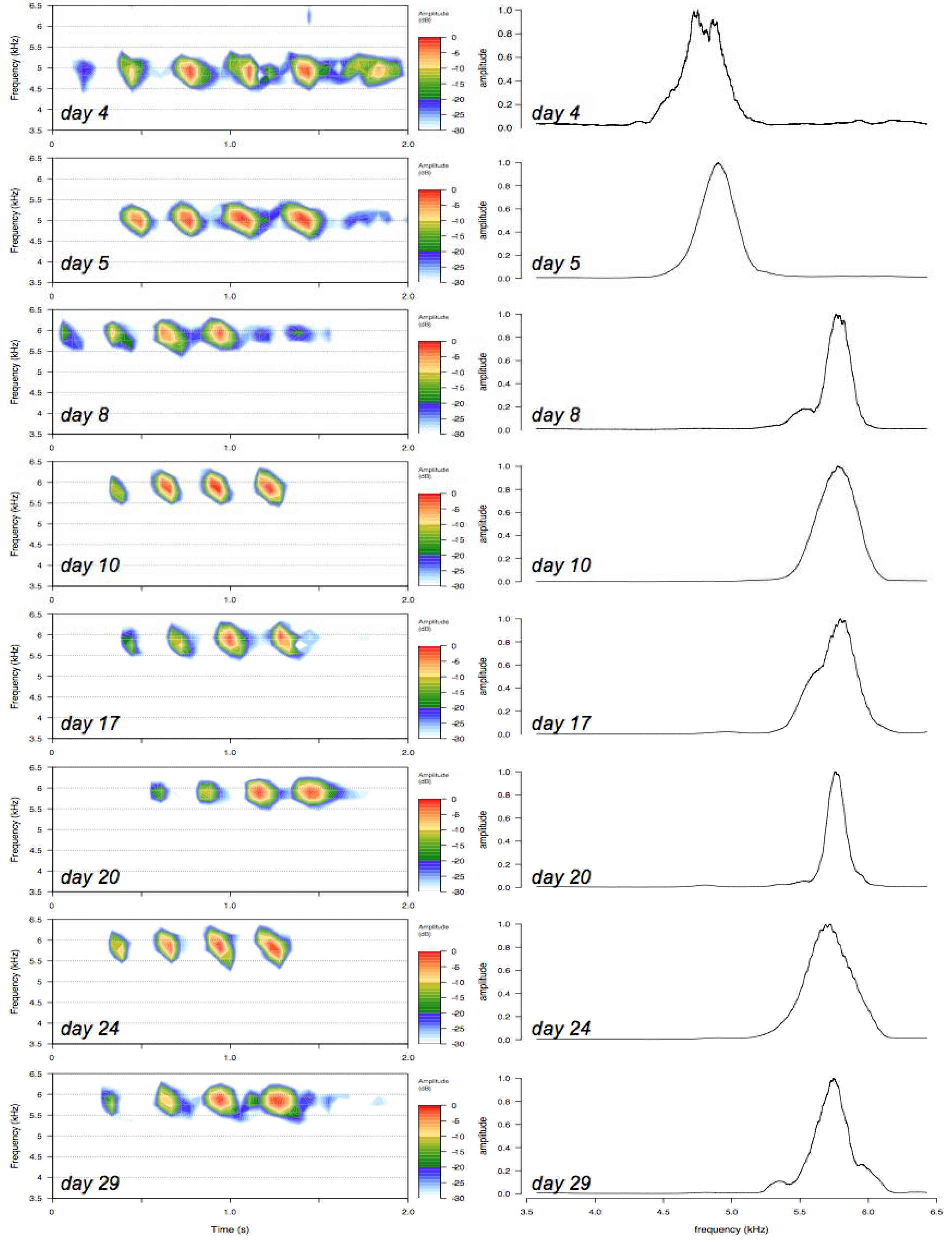
Change in peak frequency of calling songs of a representative male. Left panels are spectrograms of calling songs from a representative male that were recorded on different days. A chirp in each song are shown in this figure. The horizontal axis indicates time (seconds), and the vertical axis indicates frequency value (kHz). Color indicates relative amplitude of the frequency component as shown in scales on the right side of each chart. Figures were drawn by ‘spectro’ function in Seewave package running on R. The male ages for each chart are indicated at the lower left corner of each chart. Right panels demonstrate the distribution of frequency components. The horizontal axis shows frequency values (kHz), and the vertical axis indicates relative amplitude of each component. The amplitude values were normalized, such that the maximum values = 1.0. The date at lower left corner on each chart indicates male age. The distribution was calculated from a set of calling songs on each day (typically 10 seconds of recorded calling songs), using ‘spec’ function of Seewave package. Data were then smoothed for presentation purpose.

This observation was further confirmed at entire population level, such that there was an increasing trend in peak frequency of calling song between day 3 and day 17 (Fig. 3A). The median peak frequency of the calling songs was around 4.9 kHz on day 3, and it reached at 5.8 kHz on day 17 (Fig. 3A). Also, the variability of the frequency value showed decreasing trend between day 3 and day 17, having minimal variation around day 20 (Fig. 3B). These results demonstrate that peak frequency of the calling song converges to 5.8 kHz along with age, starting with lower and variable pitches. The final peak frequency (5.8 kHz) was comparable with the previous report (Miyashita et al., 2016a), although it should be noted that the calling songs analyzed in the previous literature was recorded between day 7 and day 14 (Miyashita et al., 2016a), and it showed slightly lower mean value (5.7 kHz), which is consistent with the observation in this study). After day 22, there was a slight downward trend in peak frequency and an increasing trend in its variability (Fig. 3A and 3B), which may be associated with senescence. However, I did not further track the calling songs after day 30 as described in the preceding section.

**Figure 3.**
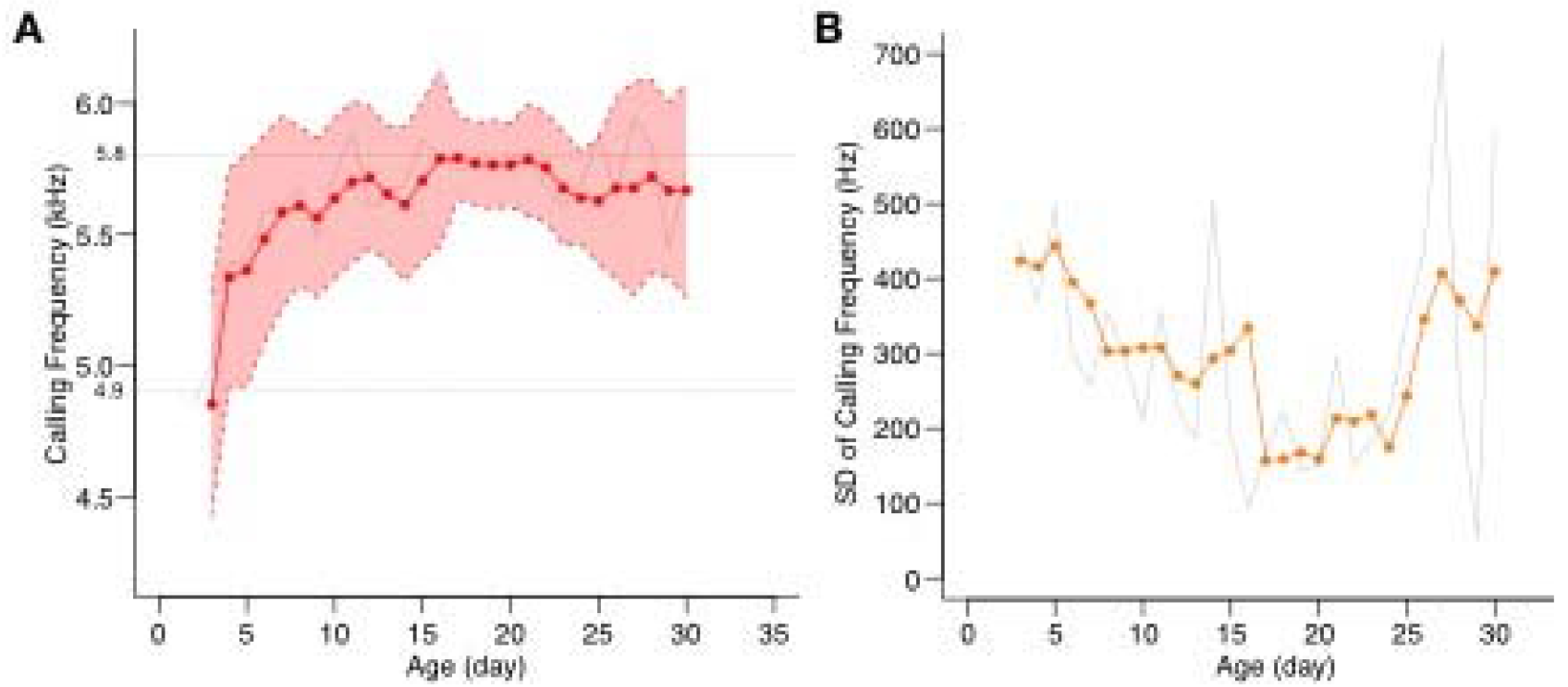
Change in peak frequency of calling songs at population level. A) A time-series data of peak frequency of cricket calling songs. This chart represents 300 recorded calling songs from 63 males. The vertical axis shows peak frequency value of calling song (kHz), and the horizontal axis shows cricket post-adult age (day of final molt = day 1). Red plots (connected by solid red lines) represents 3-day moving medians. The red-hatched region with red dashed lines indicate 3-day moving standard deviations. The number of samples (i.e. recordings) on each day is shown in Supplementary Figure S2A. The grey lines behind the red solid lines represent unsmoothed (i.e. original time-series data of median peak frequency of calling songs. A row scatterplot of entire dataset is shown in Supplementary Figure S3 for data transparency. B) Plot of standard deviation of peak frequency. The vertical axis indicates standard deviation of the peak frequency value (Hz), and the horizontal axis indicates post-adult age of crickets. The red plots connected by red solid lines represent 3-day moving medians. The original, unsmoothed data are also demonstrated in the chart with grey lines. The number of samples (i.e. recordings) on each day is shown in Supplementary Figure S2A.

### Physical explanation of the song maturation

The above finding that immature songs contained lower frequency peak indicates that the vibration frequency of the forewing membrane is lower in immature males. I assumed this was because some physical factors hindered the appropriate movement of the forewings. If this assumption is true, modifying physical parameter(s) that affects forewing movement should modify the peak frequency value of the calling songs to an appropriate value. To examine this assumption, I tested whether reducing the physical resistance of surrounding gas molecules helps crickets emit calling songs with ‘appropriate’ pitch (i.e. peak frequency at 5.8 kHz). As shown in Figure 4, the calling song of young crickets (day 5) recorded in helium substituted environment (80% Helium and 20 % Oxgen; mean molecular mass = 9.6) was significantly higher than those recorded in normal air (mean molecular mass of dry air = 29) condition (Fig. 4). The mean frequency value of the songs recorded in helium substituted condition was 5.8 kHz (Fig. 4), which was comparable with the ‘mature’ calling songs that were observed after day 17 in normal air condition (see Fig. 3A). This result supports the idea that physical resistance from surrounding gas molecules negatively impact cricket song pitch regulation, which may lead to lower and more variable frequency peak of calling songs in early adult males. Given these facts, it seems likely that mechanistic process of the song maturation in *G. bimaculatus* may involve 1) hardening (drying up) of forewings after final molt, and 2) wing muscle development which supports the fine movement of forewings. In either case, the change in peak frequency is associated with male age, which highlights a biological significance that females may use such a physical parameter of calling song as an indicator of male maturation in their reproductive behaviors.

**Fig 4.**
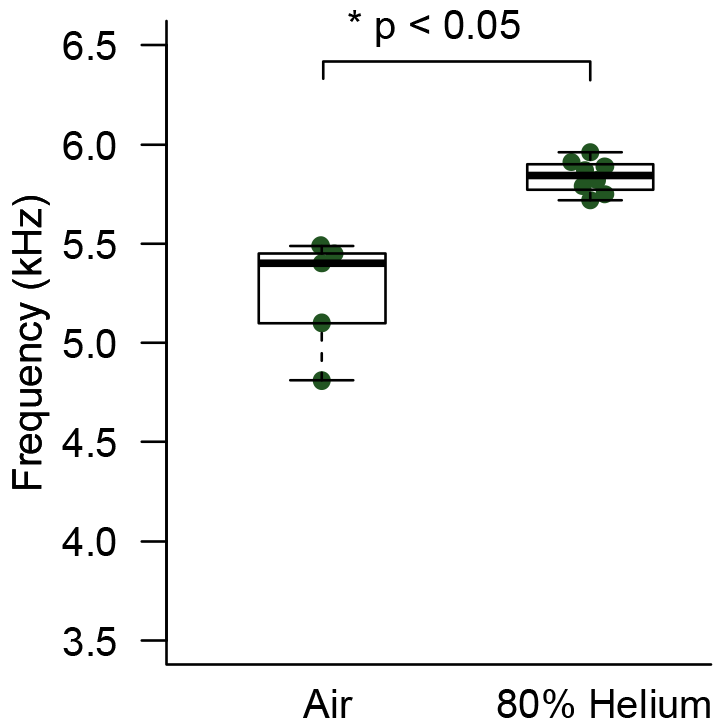
Change in peak frequency of calling songs of young males in a helium-substituted environment. The vertical axis shows the peak frequency value of calling song. The labels below the horizontal axis indicates the group: ‘Air’ represents the calling songs recorded in normal air condition, and ‘80% Helium’ represents calling songs recorded the helium-substituted condition. Males on day 5 were used for this experiment. Data from multiple different males were pooled (not all males emitted calling songs in both conditions, as sexual activity is low in early adult phase), and used for the analysis. The number of obtained samples were 5 for normal condition and 8 for helium-substituted condition. There was a significant difference of peak frequency between the two groups (p < 0.05 in Wilcoxon rank sum test).

## Discussion

The present study revealed a converging trait of male calling frequency during the post-adult maturation phase of *G. bimaculatus.* This is consistent with my previous report that the frequency values of the calling songs are robustly regulated at the late adult (7-14 days after final moult) phase (Miyashita et al., 2016b). This study provides information regarding the maturation process of the cricket mating song, in which calling song frequencies converge with male maturation. The findings further endorse the biological significance of the calling frequency value in *G. bimaculatus*.

The calling song of mature *G. bimaculatus* showed frequency peaks around 5.8 kHz, which was consistent with a previous report (Miyashita et al., 2016a). It has been suggested that female ears are tuned to particular frequency value in orthopterans (Boyd and Lewis, 1983; Katsuki and Suga, 1960; Lakes-Harlan and Scherberich, 2015; Michelsen et al., 1994). Therefore, it is likely that the change in frequency values of calling songs may provide relevant information to females for selecting mature males (but see Discussion of Miyashita et al., 2016a; the peak frequency value of *G. bimaculatus* may fluctuate depending on habitats; I have personally confirmed this between a colony in Japan and a colony purchased in United Kingdom in 2015 (Miyashita et al., unpublished data)).

Nevertheless, why do the immature males bother to sing if their songs will not be located? A possible explanation is that they ‘learn’ to sing by producing sounds at an early stage. Crickets are capable of modifying their song frequencies by auditory feedback systems (Stephen and Hartley, 1995). In this perspective, an interesting question to address in future studies is whether interference in the learning process (e.g., damaging their auditory organs) would disrupt the converging trait of the calling song.

Another possibility is that songs from immature males act as a decoy to predators. It is known that one of the ‘costs’ of cricket songs is that they increase the predation risk (Zuk et al., 2006). If the songs attract predators and only sexually mature males sing (and immature males keep silent), the predation risk should be biased towards older and sexually mature males. Also, being older means that they have survived under numbers of live threatening risks, e.g. infection risk. Therefore, it seems advantageous that young, sexually immature males bother to sing even though their songs do not attract conspecific females, because they can act as a decoy to protect more fitted and sexually mature males from predators. This point should be further investigated using both ethologic and ecologic approaches.

What molecular/physiologic/biophysical mechanism can explain the converging frequency trait? Considering the sound producing mechanism in crickets demonstrated in previous studies (Elliott and Koch, 1985; Stephen and Hartley, 1995), muscle development, tegmina condition (e.g., moisture content or tension), developmental state of the neural circuit, and other factors could affect the song frequency of the cricket. Biophysically, the parameters that determine the vibration frequency of membrane ‘instruments’ include density, tension, and size. Also, the shape of the membrane and how the initial vibration is given, etc. could affect the final frequency patterns, though it is almost impossible at the moment to establish a complete physical model. A decrease in moisture content and sclerotization of the sound source during the male maturation process may lead to a more stable vibration mode, resulting in the convergent trait of calling frequency. At least, the helium substitution experiment in this study indicates that the physical resistance from gas molecules has negative impacts to frequency stabilization, i.e. it makes song pitch lower and more variable in young immature adults. As a future study, it might be straightforward to observe vibratory properties of forewings using a laser Doppler vibrometer either in normal air or in helium substituted condition.

The study highlights the biological significance of peak frequency value of calling song in *G. bimaculatus.* In *G. bimaculatus,* a geographic divergence of the frequency value has been discussed (Miyashita et al., 2016b), and I recently further empirically confirmed it between a colony in Japan and a colony purchased in the United Kingdom (Miyashita, unpublished data). Literatures suggest that *G. bimaculatus* in European or Russian regions produce calling songs with a lower frequency than *G. bimaculatus* in Japan (Doherty, 1985; Kostarakos et al., 2009; Miyashita et al., 2016b; Popov and Shuvalov, 1977; Yamamoto et al., 2000). Whether this variation reflects different environmental selection pressures or is simply due to genetic drift remains unclear. Data from phylogenetic aspect would provide helpful information. It is worth-noting that in another cricket species, *Teleogryllus oceanicus,* mating songs are under heavy selection pressure and the songs can easily drift, or even disappear, depending on the ecologic circumstances (Zuk et al., 2006). Lastly, to fully examine behavioral significance of the calling song parameter, behavioral tests using playback sounds or synthetic sound signals is necessary. A quantitative evaluation of cricket behaviors using videotracking techniques (e.g. Perez-Escudero et al., 2014) will provide an important, robust insights.

## Acknowledgements

This study was funded by Japan Society for the Promotion of Science Research Fellowships for Young Scientists Grant #13J08664 to A.M., and by Mishima Kaiun Memorial Foundation to A.M. The funders had no role in study design. Marco Lee, Masaki Ishii and Shelley Adamo provided insightful discussions. I also thank Fumiaki Tabuchi for helping cricket survival monitoring.

